# On the relative ease of speciation with periodic gene flow

**DOI:** 10.1101/758664

**Authors:** Ethan Linck, C.J. Battey

## Abstract

Common models of speciation with gene flow consider constant migration or admixture on secondary contact, but earth’s recent climatic history suggests many populations have experienced cycles of isolation and contact over the last million years. How does this process impact the rate of speciation, and how much can we learn about its dynamics by analyzing the genomes of modern populations? Here we develop a simple model of speciation through Bateson-Dobzhansky-Muller incompatibilities in the face of periodic gene flow and validate our model with forward time simulations. We then use empirical atmospheric *CO*_2_ concentration data from the Vostok Ice Cores to simulate cycles of isolation and secondary contact in a tropical montane landscape, and ask whether they can be distinguished from a standard isolation-with-migration model by summary statistics or joint site frequency spectrum-based demographic inference. We find speciation occurs much faster under periodic than constant gene flow with equivalent effective migration rates (*Nm*). These processes can be distinguished through combinations of summary statistics or demographic inference from the site frequency spectrum, but parameter estimates appear to have little resolution beyond the most recent cycle of isolation and migration. Our results suggest speciation with periodic gene flow is a common force in generating species diversity through Pleistocene climate cycles, and highlight the limits of current inference techniques for demographic models mimicking the complexity of earth’s recent climatic history.

## Introduction

Eight decades after the modern synthesis the relative roles of gene flow, geography, and natural selection in speciation elude easy synthesis. For most of the 20th century, divergence in strict allopatry was assumed to be the dominant, if not the only, mode of speciation in most taxa, leading to an emphasis on the geography of diverging populations and an implicit suggestion of a major role for genetic drift (Dobzhansky, 1937; Mayr, 1942, 1963; Coyne and Orr, 2004; Provine, 2004). While a burgeoning body of theory has demonstrated the plausibility of speciation with gene flow in a range of circumstances (Smith, 1966; Felsenstein, 1981; Gavrilets, 2003), a lack of clear empirical examples and the related difficulty of disproving a null hypothesis of allopatric speciation did little to contest this paradigm.

More recently, a renewed emphasis on the role of natural selection in speciation in the context of adaptive radiations and “ecological” speciation has questioned the requirement of strict isolation (Schluter, 2001; Via, 2009; Nosil, 2012). The rise of genomic datasets from a wide array of model and nonmodel organisms and the concordant development of sophisticated statistical tools for demographic inference has further complicated previous assumptions (Nosil, 2008). Coalescent theory and genome-wide genetic markers provide the basis for simultaneous model selection and inference of parameter values, now implemented in a range of computational tools (Nielsen and Wakeley, 2001; Hey and Nielsen, 2007; Gutenkunst et al., 2009; Hey, 2010; Jouganous et al., 2017). As much as an early consensus from the current proliferation of studies applying these methods can be gleaned, it appears that gene flow between lineages at various points in the speciation continuum is more common than previously assumed (Mallet, 2008; Nosil, 2008; Strasburg and Rieseberg, 2008; Cui et al., 2013; Martin et al., 2013; Kumar et al., 2017; Linck et al., 2019).

While more complex models of speciation dynamics can be applied using these methods, e.g., by allowing parameter values to vary at different loci across the genome (Rougeux et al., 2017), model selection has largely treated migration (*m*; the probability a given allele is an immigrant) as an average rate in one of a maximum of two time periods (Figure 1) (Sousa and Hey, 2013). Traditional isolation with migration models are therefore distinguished from secondary contact by the presence of a time period (*T*_1_ to *T*_2_) where *m*=0. Even this relatively sophisticated model is clearly a dramatic simplification of speciation dynamics in most systems, however, where alternative assumptions of continuous gene flow (recurrent migration every generation) or strict isolation are unrealistic in most cases. For example, divergence in allopatry is widely believed to be the dominant mode of speciation in birds (Mayr, 1963; Price, 2008). Yet the extreme vagility of many bird taxa poses the question of how readily geographic classifications of speciation map on to realized rates of gene flow, leading some researchers to strongly advocate for strictly process-based definitions (Fitzpatrick et al., 2008; Mallet, 2008). Further complications arise from the imprecise, redundant language associated with the study of speciation itself (Harrison, 2012).

**Fig. 1.**
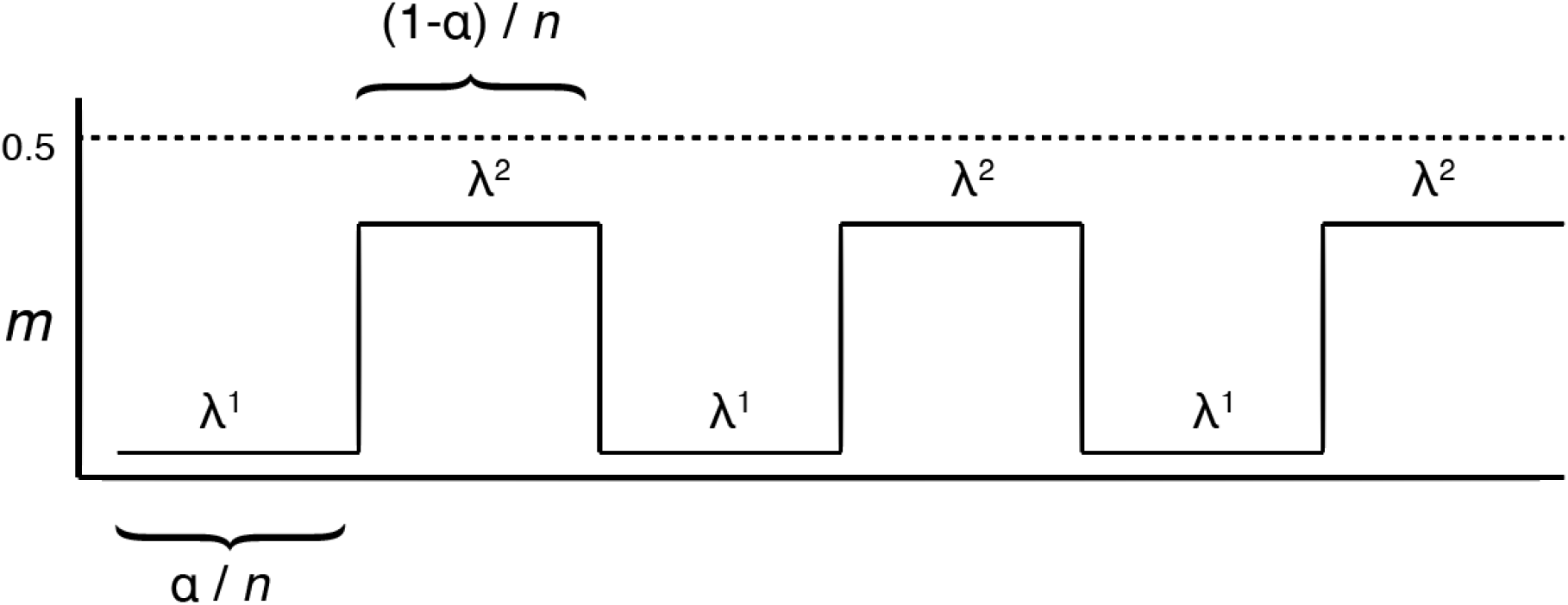
A simple model of speciation with periodic gene flow. Parameter values: *m*=probability an allele represents a migrant in a given generation; *α*= total time spent in isolation; 1 − *α*= total time spent exchanging migrants; *n*=total number of time periods; *λ*^1^= speciation rate in isolation; *λ*^2^= speciation rate with gene flow.

An alternative to either divergence in strict isolation or in the face of recurrent migration involves alternating phases of interbreeding and isolation, i.e., speciation with intermittent (or “cyclical” / “periodic”) gene flow. Cyclical processes are common over geologic, evolutionary, and ecological time scales, and include predator-prey dynamics (Elton and Nicholson, 1942), glacial cycling (Roy et al., 1996), ecological succession (Levin and Paine, 1974), and fluctuating population sizes (Vucetich et al., 1997). Yet the role of cyclical processes in generating or retarding evolutionary change has seen relatively little attention, despite evidence of unique dynamics from a handful of empirical and theoretical studies (Duckworth and Semenov, 2017; Alcala and Vuilleumier, 2014; Otto and Whitlock, 1997). Ehrendorfer (1959) proposed that “differentiation-hybridization” cycles were responsible for evolutionary patterns in the flowering plant genus Achillea, viewing hybridization as a strictly homogenizing force, an idea further developed by Rattenbury (1962) as an explanation for the diversity and persistence of the New Zealand flora. In phylogeography, Pleistocene glacial cycles in Boreal regions and the Amazon have been invoked to explain both diversification (Lovette, 2005) and patterns of genetic diversity and population genetic structure (Klicka and Zink, 1999), though the link between population subdivision and speciation remains unclear (Harvey et al., 2017).

Recently, He et al. (2019) proposed that cycles of gene flow and isolation constituted a new model of speciation, which they term mixing-isolation-mixing (or MIM), and suggest it can lead to accelerated rates of diversification. Given its relaxed requirements and flexible application to a range of potentially stochastic histories, speciation with periodic gene flow is plausibly more common than either speciation in strict allopatry or with continuous gene flow. Furthermore, recent studies on hybridization as a generative force in evolution, through either homoploid hybrid speciation (Gompert et al., 2006; Brelsford et al., 2011; Schumer et al., 2014, 2018) or adaptive introgression (Delmore et al., 2015; Norris et al., 2015; Racimo et al., 2015; Irwin et al., 2018) suggest the interaction of gene flow and isolation could promote accelerated speciation under some circumstances.

If the MIM model (hereafter “speciation with periodic gene flow”) is common and a realistic description of speciation dynamics in many circumstances, a more explicit description of its features and assessment of its consequences and genomic signature will be useful for a broad range of evolutionary biologists. As “pulse” models of admixture are widely applied in contemporary population genomics (Loh et al., 2013; Medina et al., 2018), these results may also inform our understanding of complex demographic histories below the species level. Here, we use theory and simulations to ask 1) if speciation is indeed easier to achieve under periodic migration rather than continuous migration given equivalent effective migration (*Nm*); 2) how time between migration pulses scales with waiting time to speciation; 3) whether periodic gene flow determined by a realistic scenario of glacial cycling leaves a distinct signature in commonly applied summary statistics; and 4) if joint site frequency-based approaches to demographic inference can distinguish divergence with periodic migration from traditional models of speciation with gene flow.

## Methods

### Analytical model

To evaluate the relative ease of speciation with periodic gene flow compared to speciation with continuous gene flow, we compare waiting times to speciation (average time until reproductive isolation develops between two diverging populations) using simple analytical models. First, we develop a general expectation of average waiting time given two constant speciation rates. Consider a scenario of alternating periods of migration (at a constant rate *m*>0) and isolation (*m* = 0) between populations *K*_1_ and *K*_2_, which we model as *n* alternating states of equal length *τ*, characterized by their different expected times to speciation (Figure 1). Let the rate of speciation (i.e., the probability of speciation occurring in a given time period *τ*) equal the reciprocal of the waiting time to speciation (either *T*^*a*^, meaning the waiting time to speciation in isolation, or *T*^*p*^, meaning the waiting time to speciation with recurrent migration), which we denote *λ*^1^ and *λ*^2^,respectively. The mean total expected waiting time to speciation (*T*^*t*^) is the reciprocal of the weighted sum of the speciation rates under isolation and under migration:

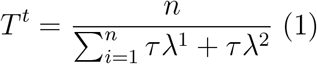

To explore the behavior of Equation 1 under different regimes of migration and isolation, we input speciation rates determined by Gavrilets’ neutral Bateson-Dobzhansky-Muller icompatibility (BDMI) model of the evolution of genetic incompatibilities on a holey adaptive landscape (Dobzhansky, 1936; Gavrilets, 2003). Full formulae and justification are provided elsewhere (Gavrilets et al., 1998; Gavrilets, 1999, 2003); a brief description of the model follows:

Consider a panmictic ancestral population of diploid organisms fixed for genotype *AABB* without recombination and then split into two completely isolated daughter populations, *K*_1_ and *K*_2_. In *K*_1_, let *A* mutate to *a* and *B* mutate to *b* unidirectionally at an equal rate of *µ* per generation. The combination of alleles *a* and *B* result in a single genotype is lethal or otherwise precludes hybrid viability; *A* and *b* remain compatible. Speciation is inevitably complete with when the genotype *aabb* is fixed in the only population with mutation (by drift alone). We define *T* as the time to speciation, or average waiting period in generations to go from the ancestral state to complete reproductive isolation. In the absence of positive selection or gene flow, i.e., a scenario of allopatric speciation by mutation and drift alone, the average waiting time to speciation will be the time to fixation of two neutral alleles (*a* and *b*), which is approximately two times the reciprocal of its mutation rate and unrelated to effective population size (Nei, 1976):

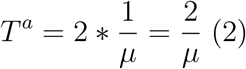

We next use Gavrilet’s (2003) extension of this model to describe parapatric speciation, i.e., speciation with limited gene flow (0<*m*<0.5) between two demes. Following subdivision of *K*_1_ and *K*_2_, a portion *m* of *K*_1_ each generation is supplied by immigrants from *K*_2_ with ancestral genotype *AABB*. Again assuming divergence in *K*_1_ is driven only by neutral mutation at a per generation rate of *µ* for both alleles, and assuming that *m* is much larger than the mutation rate, the average waiting time to speciation is given by:

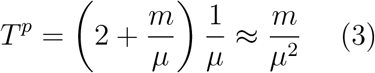

Given equal time is spent exchanging migrants as in allopatry, we calculate the arithmetic mean of these speciation rates (*λ*^*t*^):

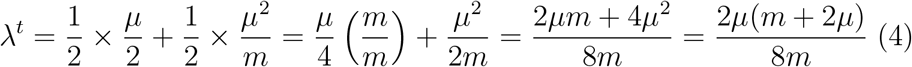

The average waiting time across all time periods is given by its reciprocal:

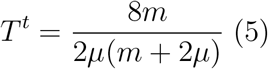

We now consider speciation with periodic gene flow if *n* time periods *τ* are unequal. Let *α* equal the proportion of time spent in isolation and 1 − *α* equal the proportion of time exchanging migrants at a constant rate *m*>0, and let the rate of speciation equal the reciprocal of the expected times to speciation *T*^*a*^ and *T*^*a*^, given by *λ*^1^ and *λ*^2^. The total expected waiting time to speciation is again simply the reciprocal of the weighted sum of the speciation rates:

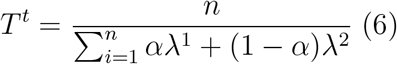

Given Equations 2 and 3, total expected waiting time to speciation can be calculated by:

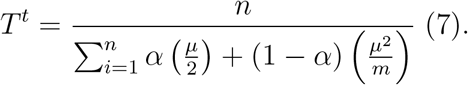

### Simulating cyclical speciation

We assessed the performance of our simple models using forward-time evolutionary simulations implemented in SLiM v. 3.1 (Haller and Messer, 2019), with diploid genotypes, constant population size, and non-overlapping generations. All parameters were chosen to closely approximate the model in Gavrilets (2003). For all simulations, we set a mutation rate of 1*x*10^*−*4^ and allowed no recombination. We modeled BDMI loci as two adjacent sites 1 bp in length; both of the genomic “regions” (BDMI locus *A* and BDMI locus *B*) on this chromosome had a specific mutation type with no fitness effect. We considered speciation complete when mutations were fixed in both BDMI loci in population *K*_2_, and did not end the simulation until this occurs; mutations were prevented from occurring on BDMI loci in population *K*_1_, which solely used as a donor of ancestral alleles in simulations involving gene flow. Outputting the final generation (i.e., waiting time to speciation) each time, we ran 100 replicates of all models. To simulate allopatric speciation, we used a single population of size *n*=100. To simulate parapatric speciation, we established two populations of size *n*=100 each, and set unidirectional migration from population *K*_1_ to population *K*_2_. We simulated speciation under periodic gene flow using three different values of *α* (the proportion of time spent in allopatry): 0.50, 0.1, and 0.01. We modified the migration rate parameter *m* across each value of *α* to fix the total effective migration (*Nm*) across the entire simulation, and initiated periods of allopatry every other generation, every 10 generations, and every 100 generations respectively. We again considered speciation complete with the fixation of both BDMI loci in *K*_1_. SLiM scripts for all simulations are available at https://github.com/elinck/speciation_periodic_migration.

### Simulating glacial cycles of isolation and contact

As a distinct goal from our analytic model of speciation and our forward time simulations to validate it, we were interested in whether cycles of gene flow and isolation between diverging populations would leave unique genomic signatures given biologically realistic parameter values. To test this, we simulated population divergence and secondary contact both 1) mediated by Pleistocene glacial cycles and 2) under a standard isolation-with-migration model using the coalescent simulator msprime v. 0.7.1 (Kelleher et al., 2016). We determined the timing and duration of gene flow between populations using empirical atmospheric *CO*_2_ concentration values from the Vostok Ice Core dataset (Petit et al., 1999). A collaborative drilling project in East Antarctica, the Vostok Ice Cores provide direct records of historical variation in atmospheric trace-gas composition through entrapped air inclusions, and cover four glacial cycles from 417,160 to 2,342 years before present (Petit et al., 1999).

We simulated a 1*x*10^7^ bp region for two populations of size *n* = 1*x*10^5^, using a recombination rate of 1*x*10^*−*8^, mutation rate 2.3*x*10^*−*9^, and allowing 10 migrants per generation (i.e., *m*=0.0001) when atmospheric *CO*_2_ fell below 250 ppm (Figure 2). This configuration effectively mimics connectivity dynamics among populations of a species distributed across a tropical montane landscape, where cooler temperatures allow the expansion of forest habitat between adjacent peaks. We started and concluded the simulation with periods of isolation, allowing 4 major periods of migration of varying duration. After completing each simulation, we calculated Wright’s *F*_*ST*_ between populations, and then used the approximation 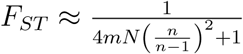(Wang, 2004) to determine a constant migration rate resulting in equivalent equilibrium *F*_*ST*_ for a paired simulation under a standard isolation with migration model (i.e., a model with continuous gene flow). We otherwise used identical parameters as our periodic migration simulation. In total, we ran 100 replicates of each scenario. We compared patterns of genetic variation between scenarios by plotting the distribution of standard summary statistics calculated both between and within populations using scikit-allel (Miles et al., 2019) implemented in Python (scripts available at https://github.com/elinck/speciation_periodic_migration). These statistics are: the number of segregating sites (SNPs); mean pairwise genetic distance (*π*); Tajima’s *D*; Watterson’s *θ*; observed heterozygosity; Weir & Cockerham’s *F*_*ST*_; *D*_*XY*_, and the average length and skew of identical-by-state haplotype blocks, a promising source of information on historical demography and dispersal (Harris and Nielsen, 2013; Ringbauer et al., 2017).

**Fig. 2.**
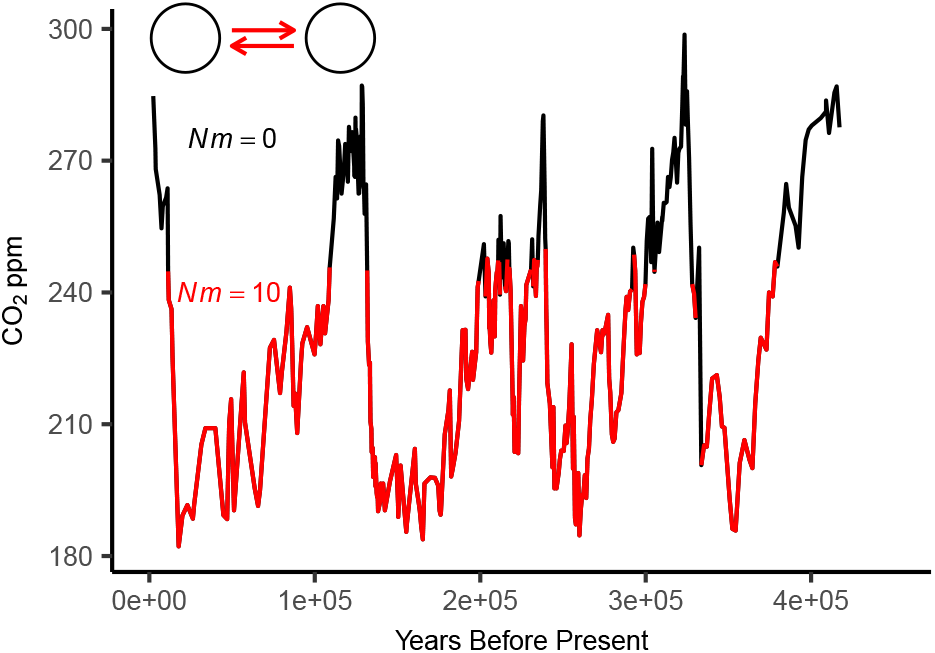
Atmospheric *CO*_2_ concentration data from the Vostok Ice Cores, color coded by migration rate in msprime simulations of populations diverging with periodic gene flow.

### Demographic inference

As speciation processes are often studied using joint site frequency spectrum-based (JSFS) demographic inference, we used our coalescent simulations to see whether this family of methods could distinguish between a model of periodic gene flow (PM) and a standard isolation-with-migration model (IM) using moments v. 1.0.0 (Jouganous et al., 2017). To model the evolution of allele frequencies under different demographic scenarios, moments analytically solves for the expected site frequency spectrum using numerical solutions of an appropriate diffusion approximation of allele frequencies (Jouganous et al., 2017). We defined two models to describe the demographic history of our simulations: a periodic migration model with four alternating periods of isolation and migration; and an isolation with migration model with continuous migration to the present (Figure 3). Both models featured symmetrical migration and independent effective population sizes (parameters *m*, *Nu*1, and *Nu*2). We ran 20 optimizations from random starting parameters for each model across 100 simulations using the “optimize log” method. For each simulation and model we used only the parameter set with the highest likelihood after optimization for downstream analysis. We then identified the model with the lowest AIC for each simulation (to correct for the higher number of parameters in periodic migration models) and examined the distribution of parameter estimates across simulations.

**Fig. 3.**
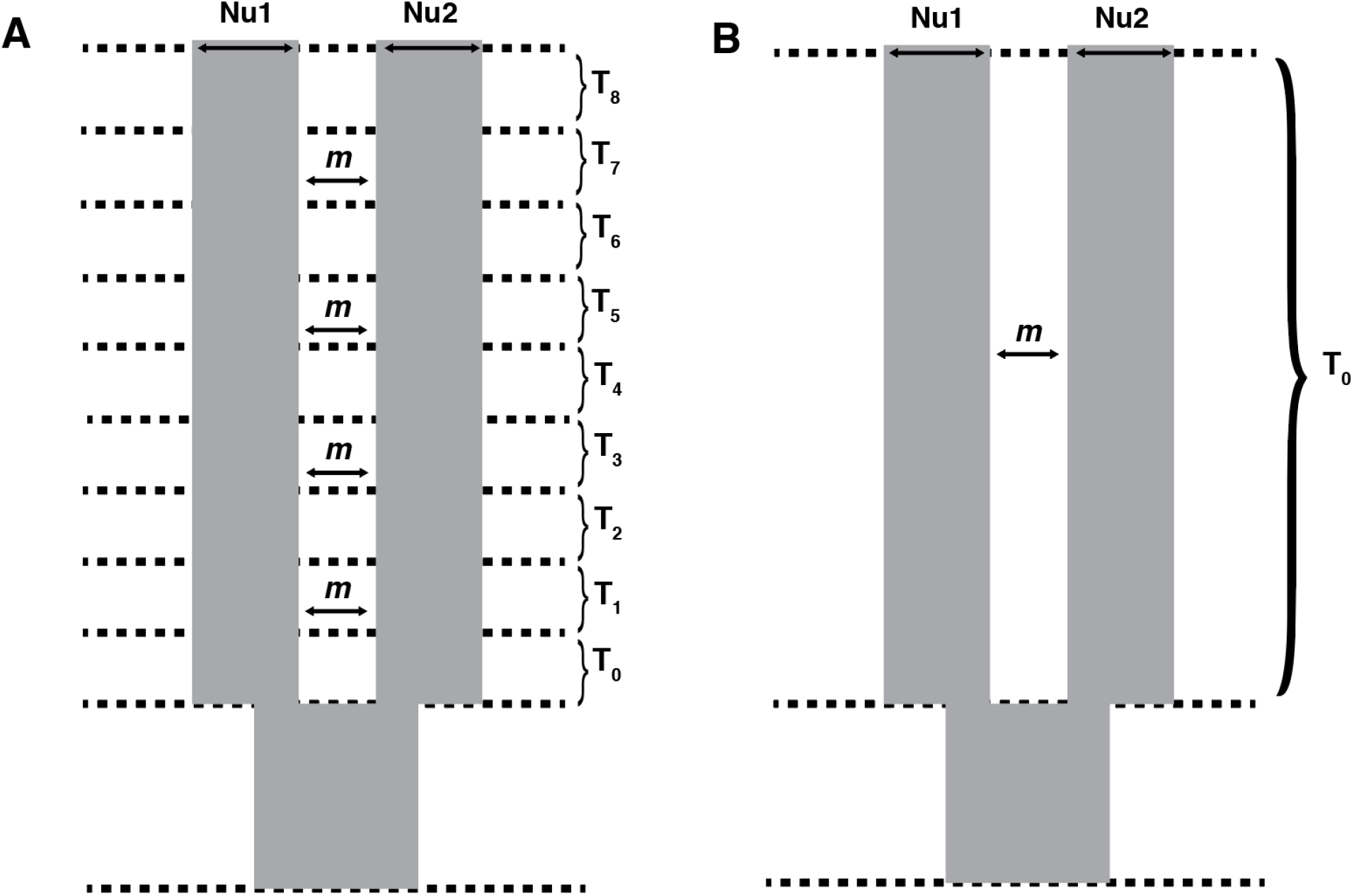
Demographic model parameters implemented in moments for A) periodic gene flow and B) constant gene flow (i.e., isolation-with-migration). *Nu*_1_ and *Nu*_2_ are parameters for the effective population size of populations 1 and 2, respectively; *m* is the symmetrical migration rate during a given time period *T*_0_ through *T*_7_.

## Results

### Analytical model

We plot *T*^*t*^ against *µ* for different values of *α* in Figure 4, generated by Gavrilet’s models of speciation by BDM incompatibilities. Though simplified, these equations capture the intuitive relationship between allopatric and parapatric speciation, e.g., that moderate gene flow erodes differences between diverging populations and results in a longer average waiting time to speciation. For example, assuming a mutation rate of *µ* = 1*x*10^*−*8^ and a migration rate of *m* = 0.1, *T*^*a*^ = 2*x*10^8^ generations under a model of allopatric speciation and *T*^*p*^ = 1*x*10^15^ in a under a model of parapatric speciation—i.e., parapatric speciation takes five million times longer. (In the absence of positive selection, speciation is clearly unlikely in both scenarios.) Notably, speciation is more likely under a model of periodic gene flow than continuous gene flow given a constant number of migrants across the entire divergence period for all three values of *α*. Further, as *α* approaches 1 − *α* in value, the average waiting time to speciation with period migration approaches its value under strict isolation.

**Fig. 4.**
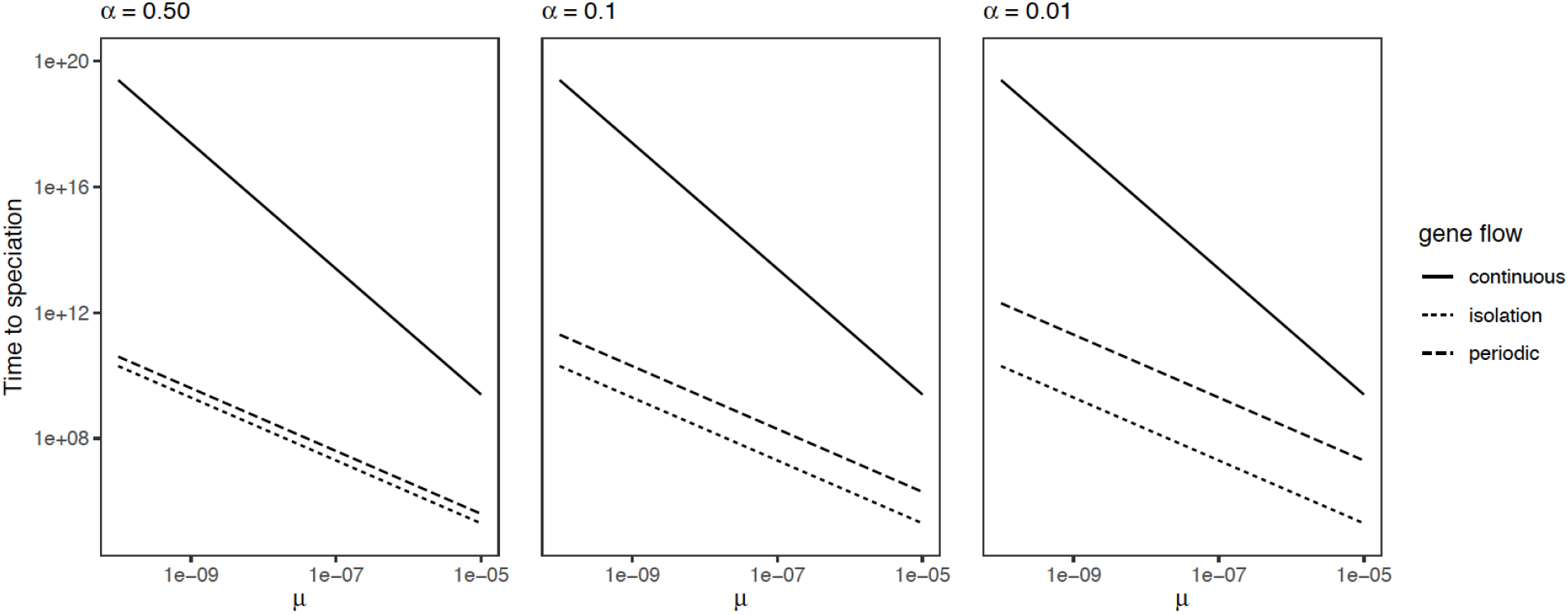
Analytical results for the relationship between BDM loci mutation rate and waiting time to speciation under alternate models of speciation, by proportion of total divergence time spent in allopatry (*α*).

### Simulating cyclical speciation

Our simulation of speciation by BDM incompatibilities broadly supported the predictions of our analytical model. The median time to speciation was lowest in isolation, highest under constant migration, and intermediate under periodic migration becoming increasingly longer as the proportion of time spent in allopatry was decreased (Figure 5). Time to speciation under periodic gene flow is bimodally distributed at all values of *α* = 1, and had the greatest interquartile range (IQR) when *α* = 0.01 and lowest when *α* = 0.01. Interestingly, speciation ocurred more rapidly than theoretical expectations in all models, with median time to speciation ranging from 18.37% to 69.99% of the expected number of generations predicted by Gavrilets (2003).

**Fig. 5.**
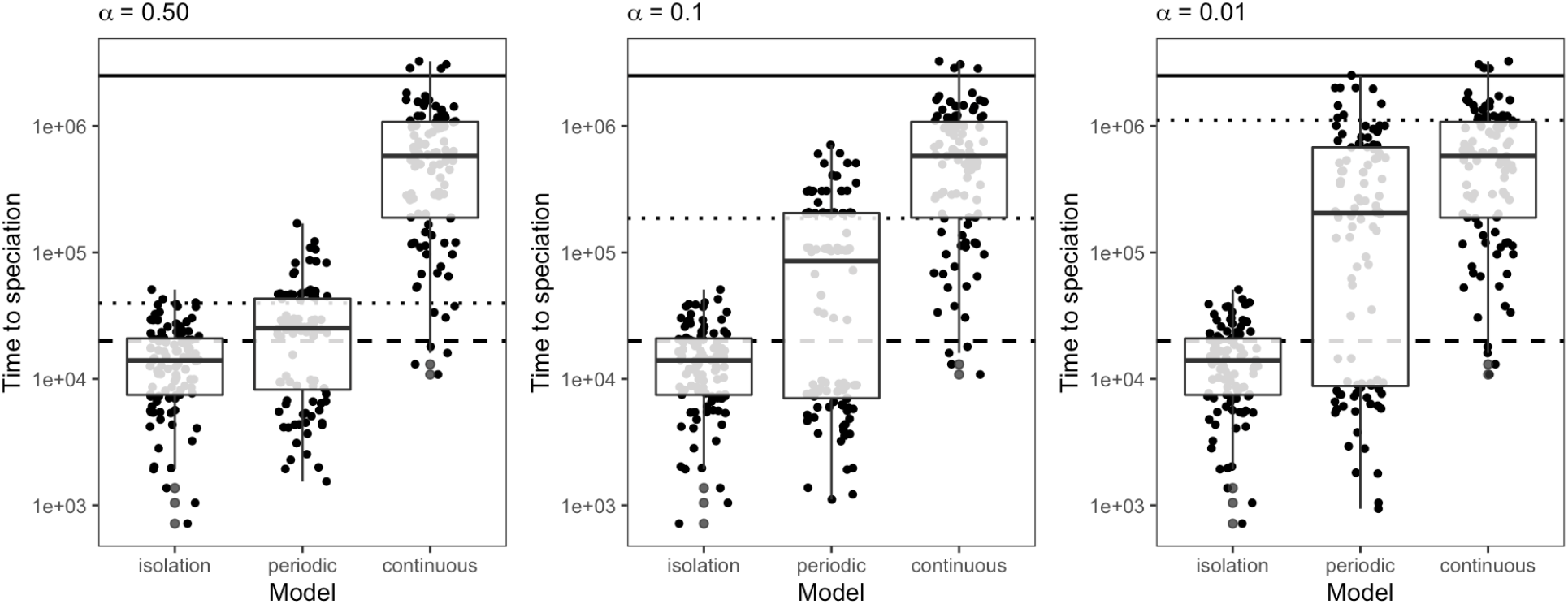
Forward time simulations of the waiting time to speciation by BDM incompatibilities under alternate models of speciation and different proportions of total divergence time spent in allopatry (*α*) Theoretical expectations of time to speciation from Gavrilets (2003) and our adapted analytical model are overlaid on the data, with the dashed line indicating time to speciation in isolation, the dotted line indicating time to speciation under periodic gene flow, and the solid line indicating time to speciation under continuous gene flow.

### Simulating glacial cycles of isolation and contact

Despite controlling for equilibrium *F*_*ST*_ (which was not significantly differed between periodic and continuous migration scenarios; Wilcox-Mann-Whitney U test, *p* = 0.89), summary statistics could easily distinguish simulations of periodic and continuous migration (Figure 6). The total number of SNPs, *π*, Watterson’s *θ*, Tajima’s *D*, observed heterozygosity, and *D*_*XY*_ were all significantly higher in continuous migration simulations (Wilcox-Mann-Whitney U test, *p* < 2.2*x*10^*−*16^). Identical-by-state haplotype blocks both between and within populations were significantly longer under periodic migration (*p* < 2.2*x*10^*−*16^). The skew of identical-by-state haplotype blocks was greater within populations under periodic migration (*p* < 2.2*x*10^*−*16^) and between populations under continuous migration, though distributions were broadly overlapping (*p* = 0.04978). There was no difference in the distribution of identical-by-state haplotype blocks over 100,000 bp in length (*p* > 0.05).

**Fig. 6.**
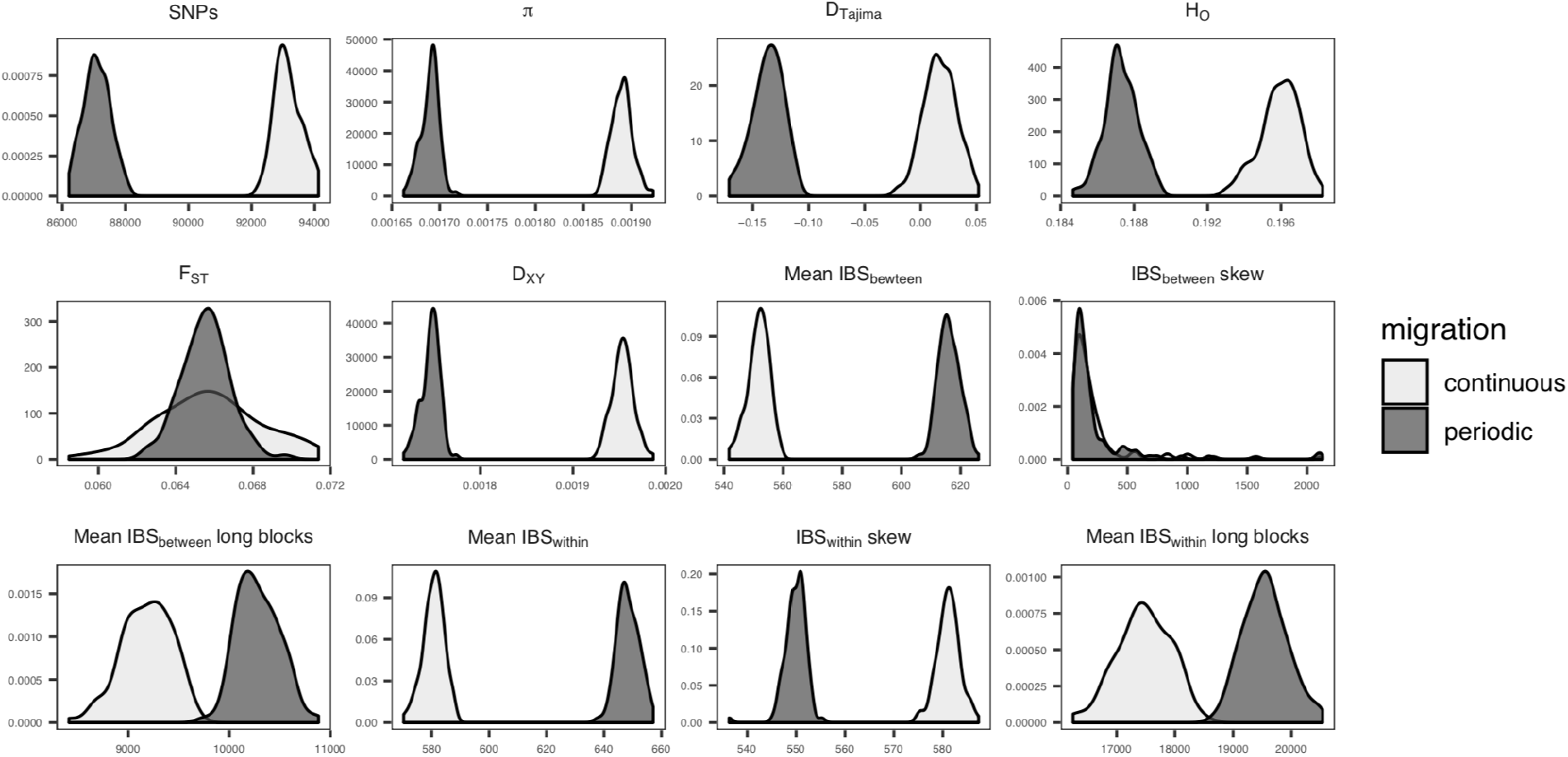
The distribution of summary statistics from simulations of periodic gene flow determined by Vostok Ice Core atmospheric *CO*_2_ concentrations (“periodic”) and a standard isolation-with-migration model (“continuous”).

### Demographic inference

Periodic migration models had lower AIC in 72.7% of simulations, with a median relative to continuous-migration models of −178.05 suggesting strong support for the periodic model (Figure 7). In most cases when continuous-migration models were preferred the corresponding periodic-migration likelihood was much lower than the median across simulations, suggesting that optimization had failed to find the global maximum. The top 5 periodic migration models accurately estimated population sizes (Figure S2) and the timing of the end of the last migration period, but appeared to have little power to estimate migration rates or the timing of earlier episodes of migration (Figure 7).

**Fig. 7.**
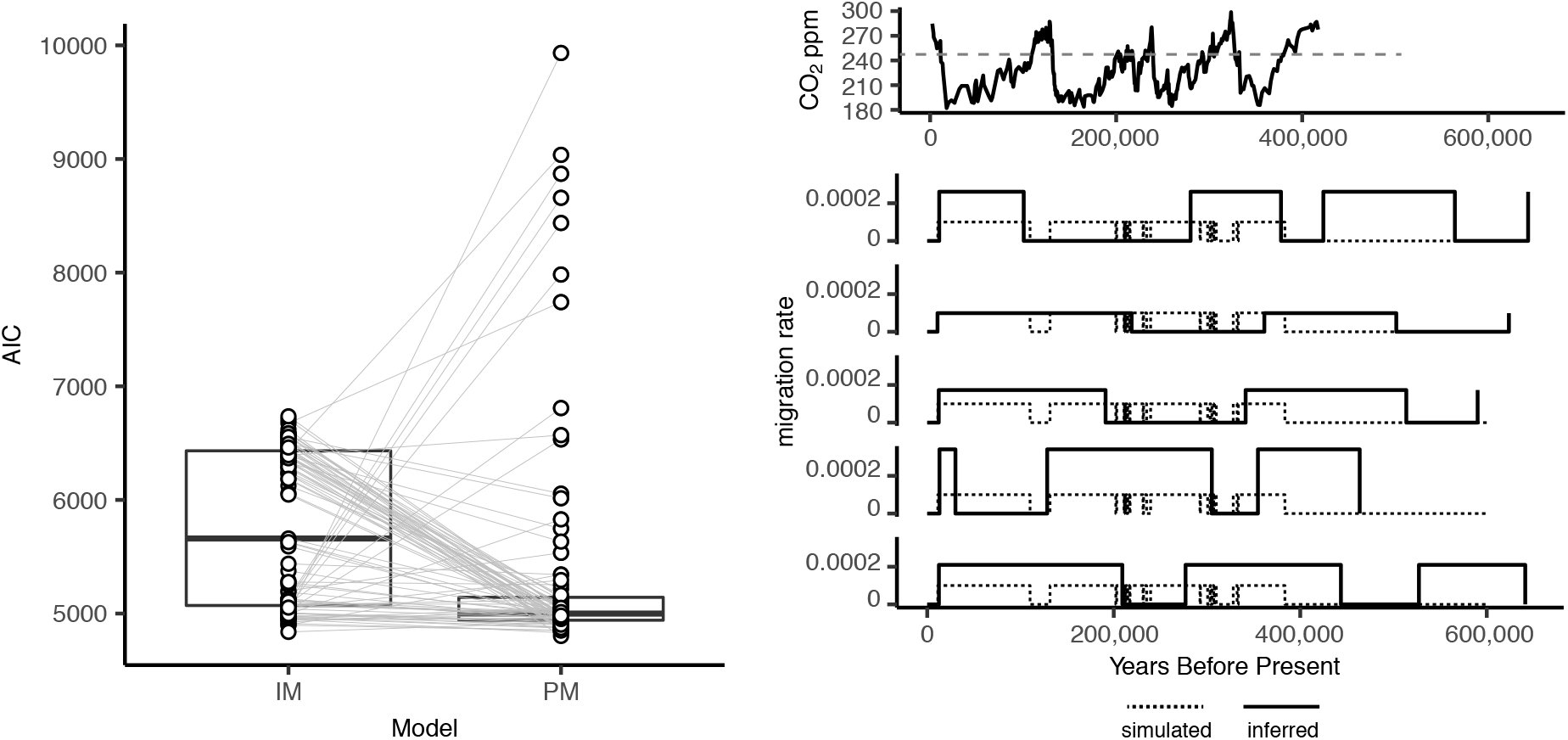
Left: AIC for continuous and periodic migration models fit to simulations with *m* = 10^*−*4^ when *CO*_2_ < 250*ppm*. Grey lines connect the maximum-likelihood parameter set for each simulation across 20 optimizations. Right: Simulated and inferred migration rates for the top 5 periodic migration model fits.

For continuous models, best-fit parameter sets were split between two regions of parameter space – high migration and ancient divergence or low migration and recent divergence (Figure S1). Parameters for the high-migration/ancient divergence regime were similar to the continuous-migration models we simulated to generate equivalent final-generation *F*_*st*_, while the low-migration/recent divergence parameters appeared to capture events since the end of the last period of migration c. 10kya. In all cases the low-migration/low-divergence-time regime had higher likelihood.

## Discussion

### Speciation with periodic gene flow

Since the inception of the field, speciation research has grappled with the question of whether reproductive isolation can develop while diverging populations exchange migrants (Darwin, 1859; Wagner, 1873; Mayr, 1963; Smith, 1966; Coyne and Orr, 2004; Fitzpatrick et al., 2008; Mallet, 2010). The inherent limitations of biogeographic definitions of speciation have lead to newer, more precise models, based on underlying processes (Schluter, 2009; Nosil, 2012) and / or the presence or absence of gene flow (Pinho and Hey, 2010). Our study of divergence with periodic gene flow found that genetic isolation is relatively more likely to occur compared to a model of continuous migration. These results suggest periodic gene flow between isolated populations may be an underappreciated force in generating species diversity under realistic conditions, and indicate gene flow-based definitions are themselves a broad category containing related but distinct speciation processes.

Our analytical model describes the expected waiting time to speciation given alternating periods of isolation and migration of specific durations and specific speciation rates (Figure 1; Equations 2 and 6). By using speciation rates determined by Gavrilets’ models of allopatric and parapatric speciation by BDM incompatibilities (Gavrilets et al., 1998; Gavrilets, 1999, 2003), we demonstrate that speciation with periodic gene flow can be nearly as likely (i.e., has a similar expected waiting time) as allopatric speciation when an equal amount of time is spent in isolation as exchanging migrants (Figure 4). Further, speciation with periodic gene flow remains substantially easier to achieve than speciation with constant migration even when as little as 1% of the total period of divergence is spent in isolation (Figure 4). This reflects an important property of our model, which is that total expected waiting time to speciation is equivalent to the harmonic mean of expected waiting times under alternate migration regimes. We thus know the average waiting time to speciation given periodic gene flow will always be less than or equal to its value given constant migration with equivalent effective migration (*Nm*). Intuitively, this application of the harmonic mean mirrors its use in estimating effective population size in the face of recurrent bottlenecks (Vucetich et al., 1997), and highlights the relationship between effective population size and effective migration rate (4*Nm*). These parameters together shape the time-averaged coalescent rate.

As it represents a specialized case of speciation in the face of periodic gene flow, we highlight several potential limitations of our simple model. First, while our validations with forward time simulations appeat to accurately reflect the expected relative median waiting time to speciation for different modes of speciation and different values of *α*, they are systematically lower than the theoretical predictions of Gavrilets (2003). This surprising result defies easy explanation, but we suspect it may result from a combination of the limitations of the Wright-Fisher model underlying our simulations and the approximations underlying the Gavrilets (2003) equations for expected waiting time to speciation. Second, we model alternating periods of isolation and gene flow as independent, “memoryless” units: partial reproductive isolation developing in one time period as one or more BDMI loci drift closer to fixation is unrelated to the probability of reproductive isolation developing in the next period, instead contributing to an average across the entire history of divergence. This might alternately lead to an underestimate (if migration substantially reduces the frequency of a BDM locus allele) or overestimate (if the frequency of the BDM locus allele is little reduced by migration after progressing substantially towards fixation) of waiting time to speciation, a result potential reflected in the bimodal distribution of time to speciation in simulations with periodic gene flow (Figure 5). Third, we ignore the possibility that reinforcement, adaptive introgression, or many other evolutionary processes might interact with modeled dynamics (though these will be discussed below). Finally, we explore only a single model for evolving reproductive isolation between diverging populations. Could other isolating barriers—e.g., the evolution of a phenotype used in mate choice—have unique behavior under periodic gene flow? We encourage others to explore this interesting question with further theory and simulations.

### Cycles of isolation and contact leave distinct genomic signatures

Repeated periods of isolation and migration between diverging populations are commonly cited in discussions of the impact of Pleistocene glacial cycles on extant biodiversity (Klicka and Zink, 1997; Avise and Walker, 1998; Klicka and Zink, 1999; Ayoub and Riechert, 2004; Lovette, 2005; He et al., 2019). Pleistocene glacial cycles and their effects on global climate (e.g., forest refugia in a drier Amazon) have been invoked as a mechanism for explaining both extant species richness and phylogeographic diversity in birds (Klicka and Zink, 1997, 1999; Weir and Schluter, 2004; Lovette, 2005; Lamb et al., 2019), plants (Vuilleumier, 1971; Bonaccorso et al., 2006; He et al., 2019), insects (Knowles, 2000; Ayoub and Riechert, 2004), fish (April et al., 2013), and mammals (Hundertmark et al., 2002). These studies have primarily evaluated a hypothesis of a Pleistocene effect by comparing molecular clock divergence time estimates between phylogroups to the timing of the last glacial maximum, sometimes to the point of controversy (Arbogast and Slowinski, 1998).

Our simulations using empirical atmospheric *CO*_2_ concentration data from the Vostok Ice Cores provide data that may be useful in more rigorously evaluating hypotheses of the role of glacial cycles. We were surprised to find that periodic gene flow was readily distinguishable from continuous gene flow by multiple summary statistics commonly applied to genomic data (Figure 6). These data suggest that neutral evolutionary processes have profoundly different effects in two scenarios that are often treated under the umbrella of “divergence with gene flow.” For example, metrics reflecting genetic diversity and effective population size (e.g., *D*_*XY*_, observed heterozygosity, *π*, Watterson’s *θ*) are reduced under periodic migration given equivalent equilibruim *F*_*ST*_, reflecting the increased strength of genetic drift during periods of isolation when the effective population size of both lineages is reduced in the absence of migration. For similar reasons, the mean length of identical-by-state haplotype blocks is greater under periodic migration: the average population-scaled recombination rate (*ρ* = 4*Ner*) is reduced across the entire timeline of divergence, preventing rapid breakdown of uniparentally inherited chromosomal regions. While our study necessarily reflects a narrow window of parameter space, we believe these statistics provide a useful starting point for detecting gene flow pulses at deeper time scales (Harris and Nielsen, 2013).

### Inferring speciation with demographic modeling

Joint site frequency spectrum based demographic model testing–conducted by comparing empirical data with the expected JSFS, as calculated by diffusion approximation (Gutenkunst et al., 2009), differential equations (Jouganous et al., 2017), or other approaches (Gronau et al., 2011)—is widely used to evaluate alternate speciation hypotheses (Hey, 2010; Nosil, 2012; Rougeux et al., 2017). While these studies have provided much of the recent evidence for speciation with gene flow (Butlin et al., 2008; Jónsson et al., 2013; Filatov et al., 2016; Rougeux et al., 2017) and led Nosil to suggest the process might be common (Nosil, 2008), they have typically employed a binary approach, contrasting models of either continuous gene flow or long periods of isolation.

Our finding that a realistic model of periodic gene flow is distinguishable from an isolation with migration model using these approaches suggests parameter-heavy models can in some circumstances be effectively applied to demographic inference. However finding the global optimum for both continuous- and periodic-migration models required starting many independent optimization runs from random starting parameters. In our analysis running 20 optimizations resulted in selecting the “correct” model in just 72% of simulations and in many cases optimization appeared to have trouble locating a global optimum. In addition, parameter estimates from these models were generally accurate for only the most recent cycle of isolation and migration. When standard IM models are fit to populations that have evolved under periodic migration, picking parameter sets with the highest likelihood tends to overestimate contemporary migration rates and underestimate divergence dates—reflecting dynamics since the end of the last period of migration rather than the full population history.

In light of these continued challenges and the reduced time to speciation under periodic gene flow compared to continuous gene flow (Figure 4), we suggest studies claiming that initial population divergence occurred in the face of gene flow argue for its plausible selective driver rather than relying on imprecise estimates of the timing of migration alone (Strasburg and Rieseberg, 2011). However, we highlight that many studies that use “speciation with gene flow” apply it broadly, including cases of secondary contact prior to the development of complete reproductive isolation (Rougeux et al., 2017). While we remain agnostic to its appropriate context, this ambiguity seems to be a continuation of a problem of language usage in speciation research more broadly (Harrison, 2012).

### Complexity in speciation and future directions

The advent of affordable genome-wide DNA sequence data for nonmodel organisms and concurrent leaps in theory and computational methods have spurred a renaissance in speciation research (Nielsen and Wakeley, 2001; Fitzpatrick et al., 2008; Mallet, 2008; Nosil, 2008; Rougeux et al., 2017). This new work has transformed our understanding of speciation, suggesting it is profoundly shaped by selection and frequently characterized by extensive introgression between diverging lineages (Mallet, 2008; Cui et al., 2013; Toews et al., 2016; Kumar et al., 2017). Our study highlights another dimension of complexity in speciation theory and research, and indicate temporal heterogeneity in rates of gene flow deserves increased scrutiny.

Beyond the findings reported in this paper, we believe several aspects of speciation with periodic gene flow provide promising research directions. First, hybridization is increasingly recognized as a generative force in evolution; adaptive introgression can increase fitness across population and species boundaries (Norris et al., 2015; Racimo et al., 2015), and occasionally lead to the evolution of reproductive isolation itself (Gompert et al., 2006; Brelsford et al., 2011; Schumer et al., 2014, 2018; Barrera-Guzmn et al., 2018). Periodic gene flow could provide raw material for these processes, increase the efficacy of selection by boosting *Ne*, or facilitate reinforcement (Coyne and Orr, 2004), accelerating speciation. Second, while the constancy and timing of gene flow between diverging populations is an important parameter and connects the historical emphasis on biogeography with the population genomic era, the selective pressures driving speciation are likely more useful for the long-term goal of developing general predictions for when and where it will occur (Kumar et al., 2017). Though difficult to identify from genomic data alone, we believe the underlying mechanisms generating reproductive isolation form the most useful framework for classifying modes of speciation, and encourage the development of new methods to identify them (Schluter, 2009). Finally, demographic inference based on the site frequency spectrum will likely remain an important tool in speciation research, but it is limited by its lack of haplotype information. Machine learning approaches approaches based on realistic forward-time simulations and multiple summaries of genetic data (e.g., Schrider and Kern (2018)) or even raw genotype matrices (Flagel et al., 2019) represent a largely untapped frontier in our quest to understand the mechanisms underlying biological diversity.

## Acknowledgements

We thank Kameron Harris for his helpful discussions on modelling. Murillo Rodrigues and Peter Ralph provided help with SLiM, and Andy Kern, Lorenz Hauser, and John Klicka provided useful feedback on the manuscript. Thanks to Nicholas Bierne for helpful comments on the first version of the preprint. This research was supported by NSF Doctoral Dissertation Improvement Grant 1701224 to J. Klicka and E.B.L, and a NDSEG Fellowship to E.B.L.

## Data availability

All code and processed data for this study are available at https://github.com/elinck/speciation_periodic_migration.

---

**Fig. S1.**
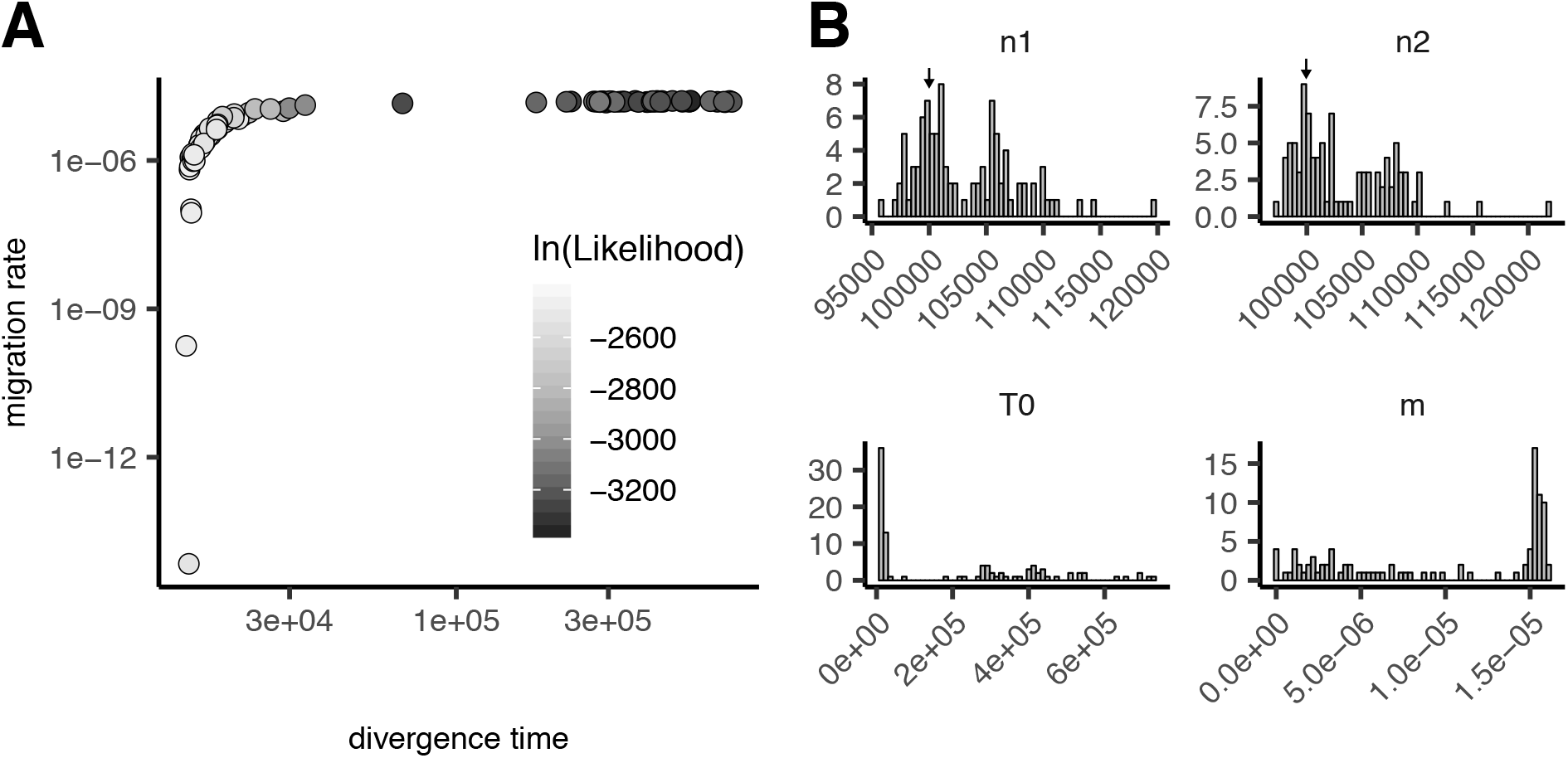
A: Best-fit parameters across 20 optimizations for each simulation. Runs with low maximum likelihoods appear to be stuck in a high-migration/high-divergence time region of parameter space. B: Summary of parameter fits for continuous-migration models fit to simulations with periodic migration. Arrows show the true population sizes.

**Fig. S2.**
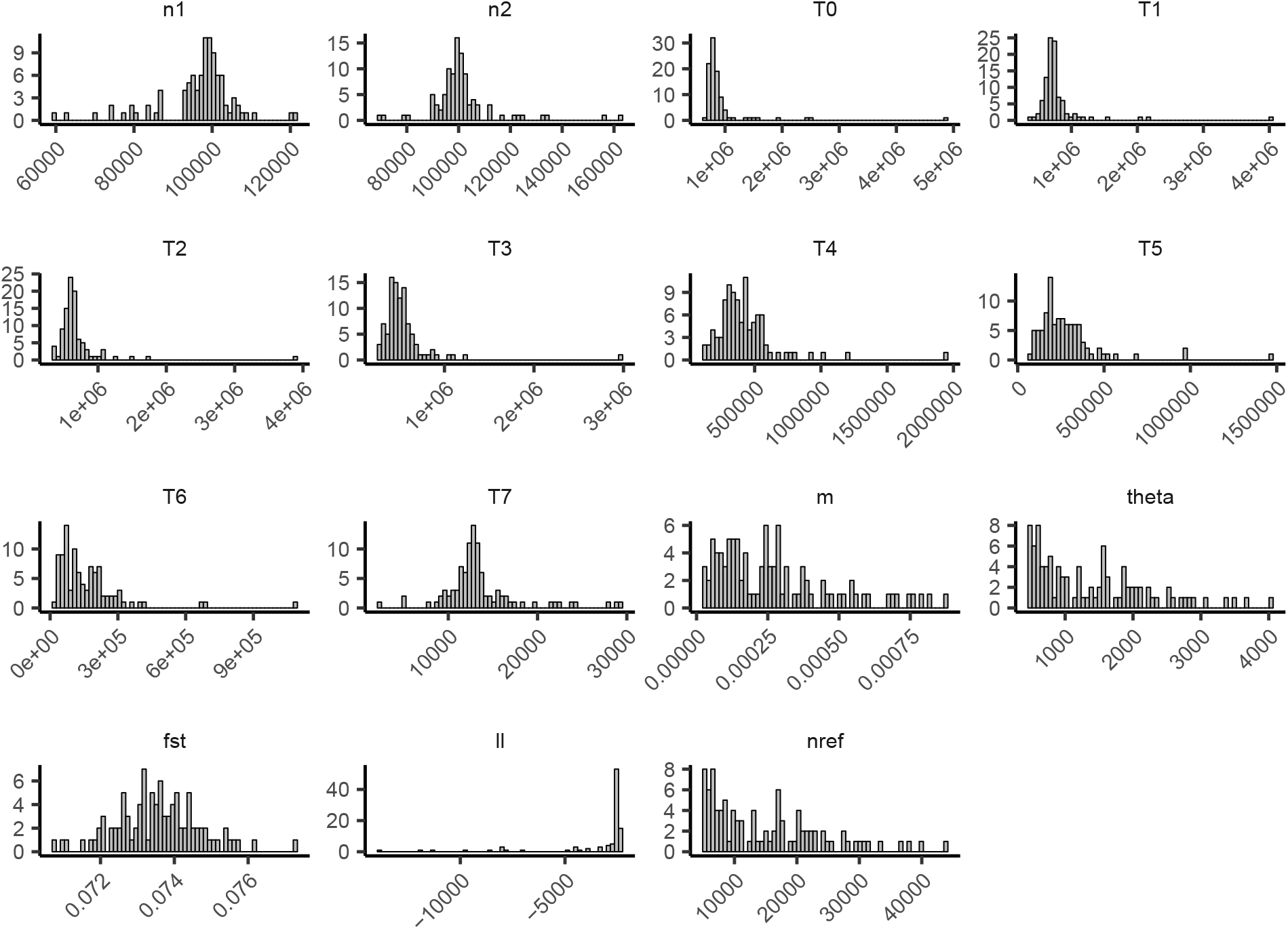
Summary of parameter fits for periodic-migration models fit to simulations with periodic migration.

